# Protein-protein complexes can undermine ultrasensitivity-dependent biological adaptation

**DOI:** 10.1101/2022.08.07.503111

**Authors:** C. Jeynes-Smith, R. P. Araujo

**Affiliations:** School of Mathematical Sciences, Queensland University of Technology, Brisbane, Australia; Institute of Health and Biomedical Innovation (IHBI), Brisbane, Australia

**Keywords:** chemical reaction networks, robust perfect adaptation, Michaelis-Menten, mass-action, ultrasensitivity, feedback

## Abstract

Robust Perfect Adaptation (RPA) is a ubiquitously-observed signalling response across all scales of biological organisation. A major class of network architectures that drive RPA in complex networks is the Opposer module – a feedback-regulated network into which specialised integral-computing ‘opposer node(s)’ are embedded. Although ultrasensitivity-generating chemical reactions have long been considered a possible mechanism for such adaptation-conferring opposer nodes, this hypothesis has relied on simplified Michaelian models, which neglect the presence of protein-protein complexes, and which are now widely acknowledged to make inaccurate predictions of signalling responses. Here we develop *complex-complete* models of interlinked covalent-modification cycles with embedded ultrasensitivity: explicitly capturing all molecular interactions and protein complexes. Strikingly, we demonstrate that the presence of protein-protein complexes thwarts the network’s capacity for RPA in any ‘free’ active protein form, conferring RPA capacity instead on the concentration of a larger protein pool consisting of two distinct forms of a single protein. Furthermore, compared to predictions by simplified models, the parametric requirements for RPA in this protein pool are much more severe, and RPA generally obtains over a narrower range of input stimuli. These surprising results raise fundamental new questions as to the biochemical requirements for adaptation-conferring Opposer modules within complex cellular networks.

## 1 Introduction

The ability to adapt to sustained disturbances and input stimuli, and to maintain key system properties within tight tolerances, is a distinctive and ubiquitously-observed feature of biological systems across all domains of life [1, 2, 3, 4, 5, 6, 7, 8]. This crucial biological response has been studied mathematically in its idealised form, known as Robust Perfect Adaptation (RPA), where the concentration of a particular molecule or activation state returns exactly to a prescribed baseline - its *setpoint* - after a disturbance or input to the system, for a wide range of possible network parameters (eg. total expression levels for the interacting molecules) [2, 4]. RPA has been identified as an essential signalling response in biological contexts as diverse as directed cell migration in single-celled organisms [9, 10, 11, 12, 13, 14, 15, 16, 17, 18, 19, 20, 5, 21], complex sensory systems [22, 23, 24, 25, 26, 27], osmoregulation in yeast and bacteria [28, 29], plasma mineral homeostasis [30], and morphogen regulation and patterning during development [31, 32]. Crucially, loss of RPA has been linked to a wide variety of diseases in multicellular organisms, including drug addiction, chronic pain, and cancer initiation and progression [33, 34, 35, 36, 37, 38].

Importantly, RPA is a structural property of networks, and is not dependent on fine-tuning of system parameters. The complete solution space for RPA-capable network designs has now been identified in full generality [4], for networks of unlimited size and complexity. In fact, it is now known that all RPA-capable networks are decomposable into modules, of which there are two broad, yet well-defined, classes: Opposer modules, and Balancer modules [4]. Opposer modules are a generalisation of the three-node RPA solution known as ‘negative feedback with buffer node’ (NFB) first identified by Ma et al. [8] through extensive computational searching. All Opposer modules contain at least one negative feedback loop, and have at least one computational node (known as an ‘opposer node’) embedded into the feedback loop. In complex networks consisting of a large number of interacting molecules, Opposer modules may contain distributed integral controllers known as ‘Opposing Sets’, consisting of multiple interlinked negative feedback loops containing special arrangements of opposer nodes [4]. Balancer modules, by contrast, generalise the three-node ‘incoherent feed-forward loops with proportioner node’ (IFFL) identified by Ma et al. [8]. Balancer modules consist of an arbitrary number of parallel pathways, at least two of which must be incoherent in nature, and which have special computational nodes known as ‘balancer nodes’ embedded into them [4]. In general, RPA-capable networks can contain any number of Opposer and/or Balancer modules, connected together according to well-defined interconnectivity rules, to orchestrate highly complex RPA-capable networks [4, 39].

A central question in biochemical network theory remains how complex organisms incorporate these well-defined RPA-conferring network structures into their signalling networks [4, 39]. Until now, most known Balancer modules have been identified in very small signalling networks such as two-component regulatory systems in bacteria [29, 40], and two- or three-node transcription networks across a variety of cell types [41, 42]. By contrast, RPA in large and highly complex signalling networks such as mammalian signal transduction networks is thought to depend on feedback structures, and hence, Opposer modules [1, 4, 39].

The essential ingredient for all Opposer modules is the embedding of at least one *opposer node* - a collection of chemical reactions which computes the integral of the deviation between the RPA setpoint and the instantaneous signalling level of an RPA molecule – into the overarching feedback structure of the module [4]. Currently, there are just two known types of opposer node - antithetic integral control [43, 44], and ultrasensitivity-generating motifs [45, 39]. Antithetic integral control is a universal ‘opposer node’ that confers an exact (i.e. ‘perfect’) form of RPA, which has been identified in endogenous cellular networks in the form of sigma/anti-sigma factors [40], for instance. Antithetic integral control has also been implemented successfully in a variety of synthetic bionetworks [2, 44, 43, 21]. Ultrasensitivity-dependent opposer nodes, on the other hand, remain a theoretical class of RPA-promoting signalling structures that are thought to confer an ‘almost perfect’ form of RPA [8, 39, 45, 46], and although they are posited to play a role in the regulation of cellular signal transduction pathways [1], specific instances of this type of opposer node are yet to be identified in cells. We depict the relationship between ultrasensitivity and RPA in Fig 1.

**Figure 1:**
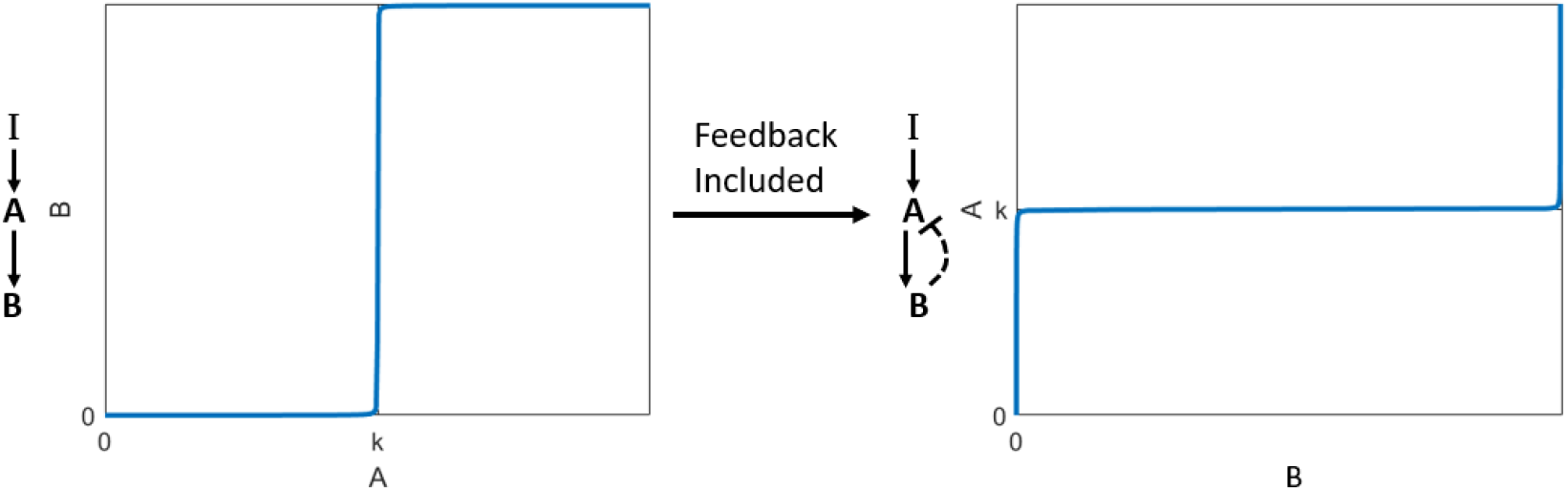
The relationship between ultrasensitivity and RPA. In an open-loop signalling cascade (left), an ultrasensitivity-generating mechanism at the level of a single protein (*B*) creates a reversible switch whereby the output can exist in either a low activity state, or a high activity state, with a vanishingly narrow transition zone (for *A* ≈ *k*) between the two states. If this ultrasensitivity-generating mechanism is embedded into a negative feedback loop (right), the narrow transition zone for the ultrasensitive switch is converted into a narrow tolerance around an RPA setpoint (*k*) for the upstream molecule *A*.

The mathematical evidence for the creation of RPA-promoting opposer nodes via ultra-sensitive switches has largely stemmed from the landmark computational study undertaken by Ma et al. [8]. We depict the essential mathematical framework of the three-node signalling structures considered by Ma et al. [8] in Fig 2(a). In that study, the focus is specifically on three nodes networks and therefore the feedback structure was predicated on the input and output nodes being distinct from one another. It was later shown [1, 4, 39] that two nodes are sufficient for RPA in a feedback structure (Opposer module), since input and output nodes need not be distinct as shown in Fig 2(b). The key feature of these RPA-conferring models by Ma et al. [8] was the embedding of an ultrasensitive switch at the location noted as an ‘opposer cycle’ (referred to as a ‘buffer node’ in the study by Ma et al. [8], which was restricted to three-node networks).

**Figure 2:**
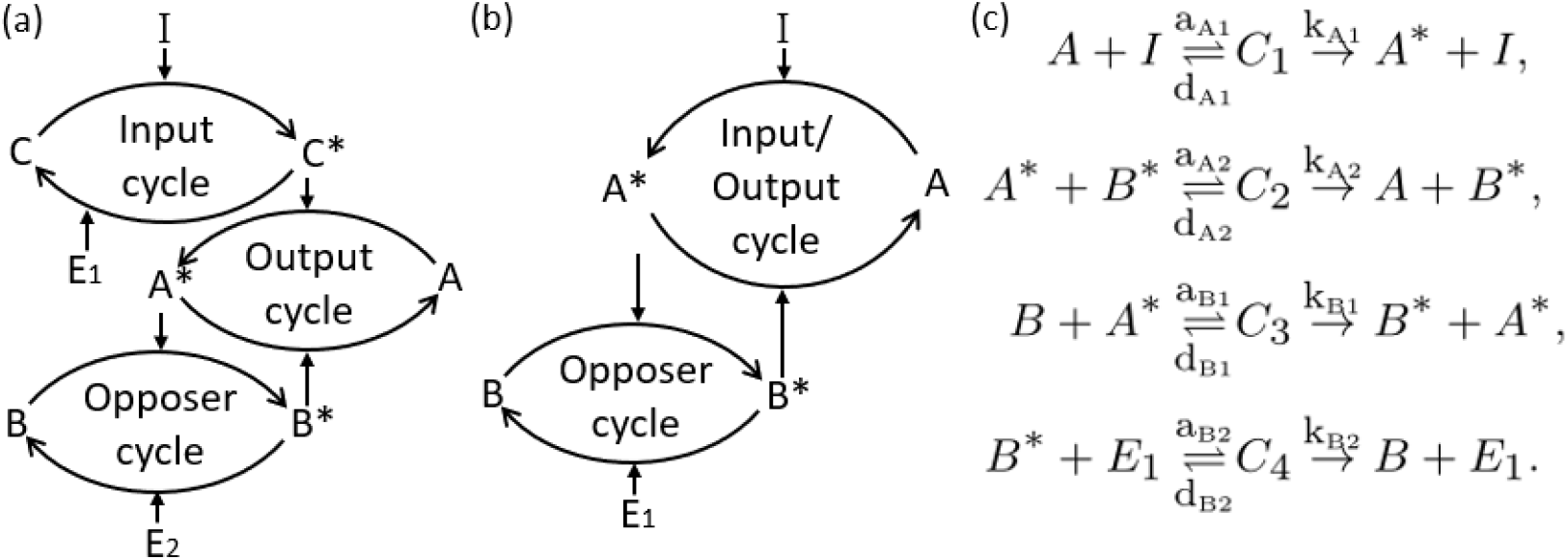
Network diagrams for (a) the three-node network analysed by Ma et al. [8], and (b) the corresponding reduced two-node network with a single node representing both input and output. (c) The graph structure for the set of chemical reactions corresponding to the network in (b), from which either *complex-complete* mass action equations [45] or simplified Michaelis-Menten equations (neglecting all protein-protein complexes) can be derived. In each case, the active form of protein *X* is denoted *X*^*∗*^. The four intermediate protein-protein complexes are denoted by *C*_1_, *C*_2_, *C*_3_, and *C*_4_. The association, dissociation, and catalytic rate constants are given by by *a*_*i*_, *d*_*i*_, and *k*_*i*_, respectively, as shown.

Importantly, Ma et al. [8] drew conclusions as to the putative RPA-capacity of this network under the simplifying assumption of Michaelis-Menten kinetics for each covalent-modification cycle, Eqs (1)-(2), which pre-supposes that the concentrations of any intermediate enzyme-substrate complexes may be neglected - an assumption which is generally thought to be reasonable for a single covalent-modification cycle in isolation when the total concentrations of the interconverting enzymes (e.g. *I*_*tot*_, *E*_*tot*_) are exceedingly small in comparison with that of their substrates (*A*_*tot*_, *B*_*tot*_) [47]. In many complex signalling networks of biological interest, such as cancer signal transduction networks, enzymes and substrates typically exist at comparable concentrations [50, 51, 52, 53, 54], and covalent-modification cycles are highly interlinked [45].

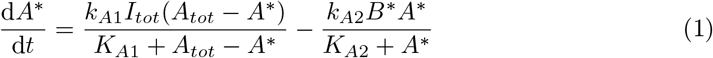

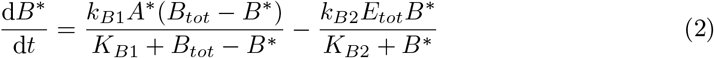

Ma et al. [8] demonstrated analytically that RPA is dependent on the Michaelis constants in the opposer cycle (*K*_*B*1_, *K*_*B*2_) being very small in comparison with the corresponding substrate abundance (*B*_*tot*_ *− B*^*∗*^, and *B*^*∗*^, respectively) (Fig 3(a)). This creates a (zero-order) ultrasensitive switch in the opposer cycle [48]. The RPA setpoint, *σ*, can then be derived from Eq (2) at steady state, where

**Figure 3:**
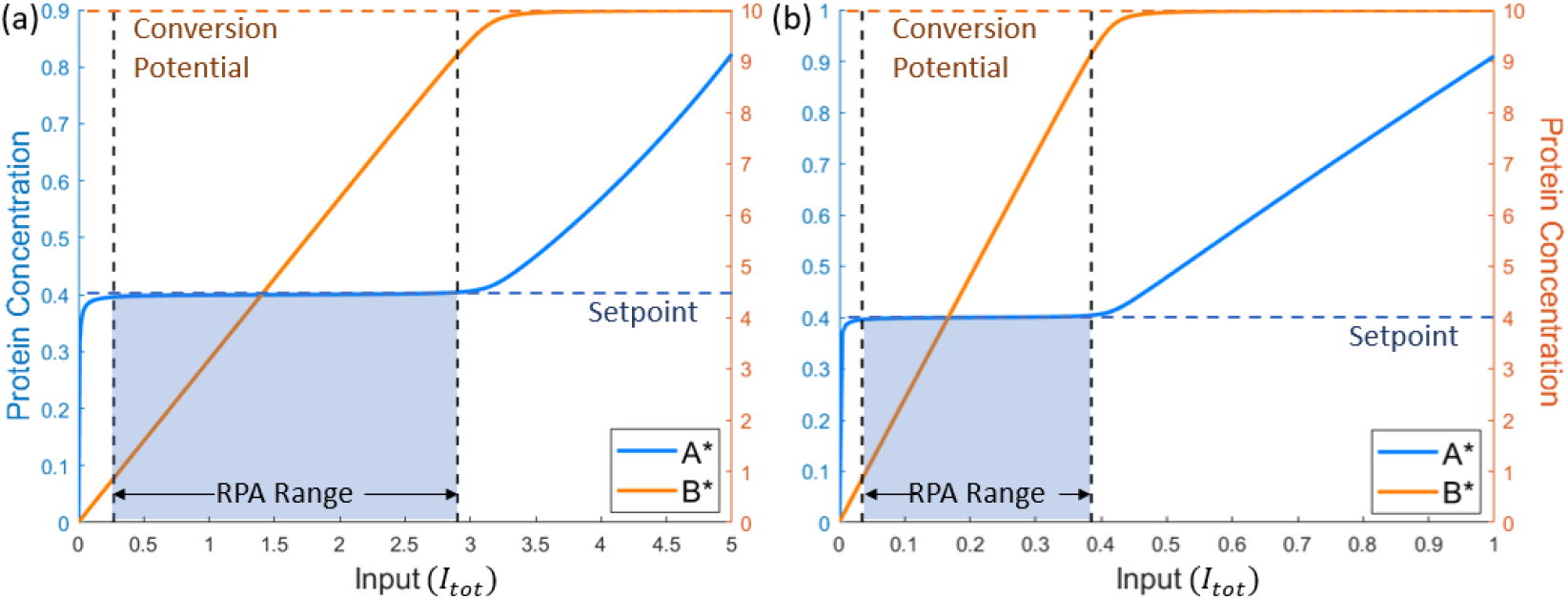
Example simulations of (a) Eqs (1)-(2) and (b) Eqs (4)-(5), employing parameter sets used by Ma et al. [8], and Ferrell [1], respectively. We highlight the range of input values over which the RPA property can be observed i.e. the ‘RPA Range’. The beginning of this range is determined by the value of input when *A*^*∗*^ is first within 1% of the estimated setpoint (dark blue dashed line), and ends when *A*^*∗*^ deviates from the setpoint by more than 1% and *B*^*∗*^ reaches the conversion potential (dark orange dashed line). Parameters: *A*_*tot*_ = *B*_*tot*_ = 10, *E*_*tot*_ = 1, *k*_*A*1_ = *k*_*A*2_ = 200, *k*_*B*1_ = 10, *k*_*B*2_ = 4, *K*_*A*1_ = *K*_*A*2_ = 1, and *K*_*B*1_ = *K*_*B*2_ = 0.01.

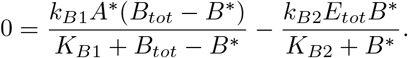

In the limit as *K*_*B*1_, *K*_*B*2_ → 0, the steady-state condition tends to

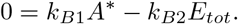

The estimated setpoint (for the output *A*^*∗*^), is therefore given by

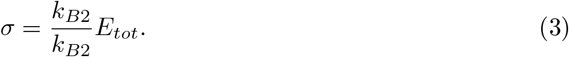

Thus, *σ* is the basal level to which the output returns following any persistent disturbance to the network (*I*), and the location of the ultrasensitive switch in the ‘open-loop’ cascade (see Fig 1).

For the sake of a complete discussion of simplified RPA frameworks, we note that Ferrell [1] later considered an even simpler version of the two-protein version of the Ma et al. [8] model, using simplified mass-action kinetics rather than Michaelian kinetics for the input/output cycle (Eq (4)). We depict a characteristic output of this system in Fig 3(b) using an identical parameter regime as the corresponding RPA-promoting Michaelian model (Fig 3(a)). As shown in Fig 3, the qualitative responses are almost identical, with a variation in the observed ‘RPA range’ being the only difference between the outputs for the two model variations. The Ferrell model [1] most closely resembles the embedding of a covalent modification cycle into a post-transcriptionally orchestrated feedback loop, requiring de-novo protein synthesis. By contrast, the Ma et al. model [8], involving two interlinked covalent modification cycles, corresponds to short-term signalling within signal transduction cascades.

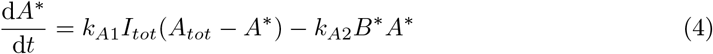

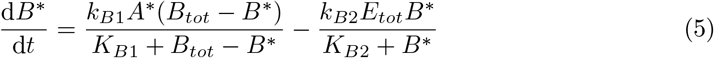

The pitfalls and potential inaccuracies of employing the simplified Michaelis-Menten equations for the modelling of signal processing through covalent-modification cycles are now widely appreciated [45, 46, 49]. Alternative quasi steady-state assumptions (QSSAs) have been successfully employed to obviate some of these difficulties in relatively simple models [46, 55, 56], but these approaches elude covalent modification cycles with added regulations such as positive autoregulation (PAR) [45], or the intricate interconnection of multiple linked covalent-modification cycles. As a consequence, it is currently unknown to what extent ultrasensitivity-dependent opposer nodes could exist in biology - either endogenously within vast and complex signal transduction networks, or in a synthetic setting.

In this paper we ask a fundamental question: has the prevalent use of simplified models given a false hope for ultrasensitivity-driven RPA mechanisms? In ‘real’ enzyme-mediated signalling networks, enzymes can only exert their activities through binding events with their substrates, creating enzyme-substrate complexes which could exist at non-negligible concentrations. Can RPA still obtain when the presence of all of these protein-protein complexes are considered?

## 2 Methods

We construct here a minimal version of an ultrasensitivity-driven Opposer module, which explicitly considers all protein-protein interactions, and all intermediate molecular species - a framework we call *complex-complete* [45]. First, we consider the mass-action equations induced by the reaction scheme depicted in Fig 2(c):

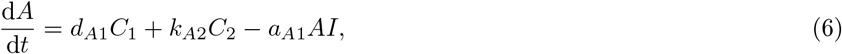

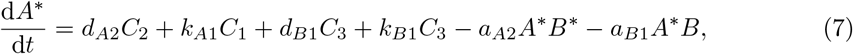

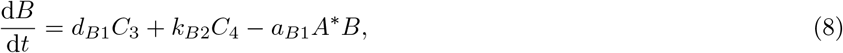

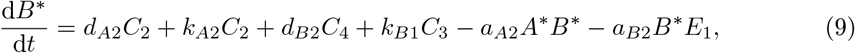

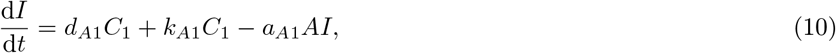

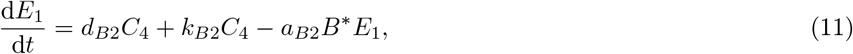

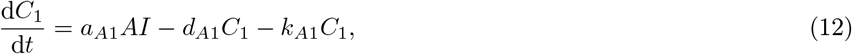

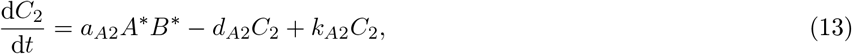

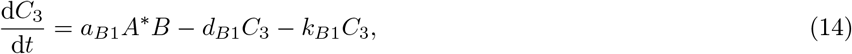

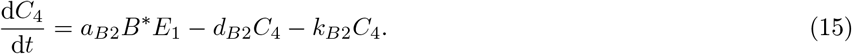

Taken together, these reaction rates reveal the following four mass conservation relations:

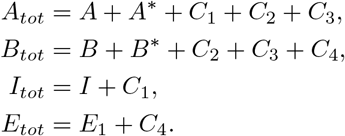

In this study we examined 10^5^ parameter sets whereby individual parameters were selected to be of the form 10^*n*^, with *n* a uniformly distributed real number on the interval [−3, 4] for the catalytic constants and Michaelis constants, and on the interval [0, 4] for total protein abundances. We computed protein steady-states by simulating the system of mass-action equations (6)-(15) using Matlab’s ODE solver, ‘ode23s’. This solver avoids timescale issues that can occur in models with parameters and variables of largely varying magnitude [59, 60, 61]. We provide all our code in an online repository (see Data Availability statement).

For each parameter choice, we compute a steady-state dose-response profile, recognising that RPA is characterised by two essential features: (i) The output steady-state must ‘track’ the setpoint (predicted to be 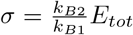, see Eq (3)), for a non-negligible range of inputs to the network. This range of inputs is referred to hereafter as the *RPA range*. (ii) The steady-state concentration of some *other* protein (in this case, some form of *B*) must vary as the input varies, across the RPA range. This second condition is essential to distinguishing ‘true’ RPA from any ‘trivial’ form of RPA that can occur when any protein concentration reaches its maximum possible value. We call this maximum possible value its *conversion potential* [45] (see Fig 3). It is now well-established that all RPA-capable networks have at least one ‘non-RPA’ variable, which acts as an ‘actuator’ node [40].

To determine the RPA Range, we identify the smallest (*I*_*S*_) and largest (*I*_*F*_) values of the network input for which RPA obtains (if at all); then *range* = *I*_*F*_ − *I*_*S*_. Thus, *I*_*S*_ is determined by identifying the smallest input value for which (i) the RPA-exhibiting variable, at steady state, is unchanging for successive input values (within a tolerance of 2%), and is within a small tolerance (±1%) of the estimated setpoint, and (ii) some other (non-RPA) variable is changing, at steady state, by at least 1% across the same successive input values. Then, *I*_*F*_ (>*I*_*S*_) is identified as the input value for which these conditions are first violated.

For every parameter choice, we determine whether the network is capable of achieving the RPA property by identifying values for *I*_*S*_ and *I*_*F*_, if they exist. We then partition the parameter regimes into two groups based on whether the associated network is able to achieve RPA or not. Where RPA is achieved, we further partition the parameter regimes into subgroups based on the magnitude of the RPA range.

On the basis of these comprehensive parameter searches, we also present in the next section some more ‘focussed’ results, for illustrative purposes, highlighting the roles of key parameter groups (Michaelis constants, catalytic constants, total protein abundances, etc) and their influence on RPA capacity and/or of the RPA range, while holding all other parameters fixed.

## 3 Results

We find that the RPA-capacity of the complex-complete model of embedded ultrasensitivity exhibits a number of unexpected characteristics, deviating in at least three fundamentally important ways from the predictions of simplified (Michaelian) models of ultrasensitivity-driven RPA. We consider each of these surprising findings in turn below.

### 3.1 RPA is *never* achieved in the ‘free’ active form of the regulated protein (*A*^*∗*^)

In the Ma et al. study [8], employing Michaelis-Menten kinetics for all enzyme catalysed reactions, it was clear that the active form of the ‘output’ protein, *A*^*∗*^, could exhibit RPA for a wide range of parameters and system inputs provided that a protein in the feed-back portion of the circuit (i.e. the opposer protein, *B*^*∗*^, in our nomenclature) exhibited ultrasensitivity (*K*_*B*1_, *K*_*B*2_ ≪ *B*_*tot*_). When we relax the Michaelian assumption of negligible enzyme-substrate complexes, however, and track all protein species explicitly via our complex-complete framework, we find that *A*^*∗*^ - the ‘free’ active form of the output protein - *never* tracks the estimated setpoint (*σ*), or any other non-trivial setpoint, for any choice of biochemical rate constants or protein abundances.

Instead, we find that RPA can be achieved by an entirely different biochemical species. In particular, *C*_3_, being the enzyme-substrate complex consisting of the *active* form of *A* (enzyme) bound to the *inactive* form of *B* (substrate), can achieve a concentration equal to *σ*, for a significant range of inputs, and for a wide variety of parameter choices. Intriguingly, under conditions where the concentration of *C*_3_ was found to track the estimated setpoint, we discovered that the concentration of *A*^*∗*^ was identically zero (or, at least, less than our chosen tolerance - see Methods). From this, we conclude that the ‘true’ RPA variable for a feedback system with embedded ultrasensitivity is the ‘total’ input to the opposer cycle which, in this case, is given by 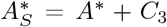. Indeed, 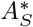 corresponds to the location of the ultrasensitive switch in the Goldbeter-Koshland model of zero-order ultrasensitivity [48, 45, 62]. As we show in Fig 4, when 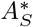 tracks the value *σ* (Fig 4(a)), *A*^*∗*^ = 0 (Fig 4(b)). At the same time, *C*_3_ also tracks the value *σ* (Fig 4(c)). In addition, *C*_4_ = *E*_*tot*_ (Fig 4(d)), and *E*_1_ (the concentration of free enzyme) is identically zero (not shown). We recognise that these conditions correspond to the complete saturation of both interconverting enzymes for the opposer cycle - a hallmark of zero-order ultrasensitivity [48].

**Figure 4:**
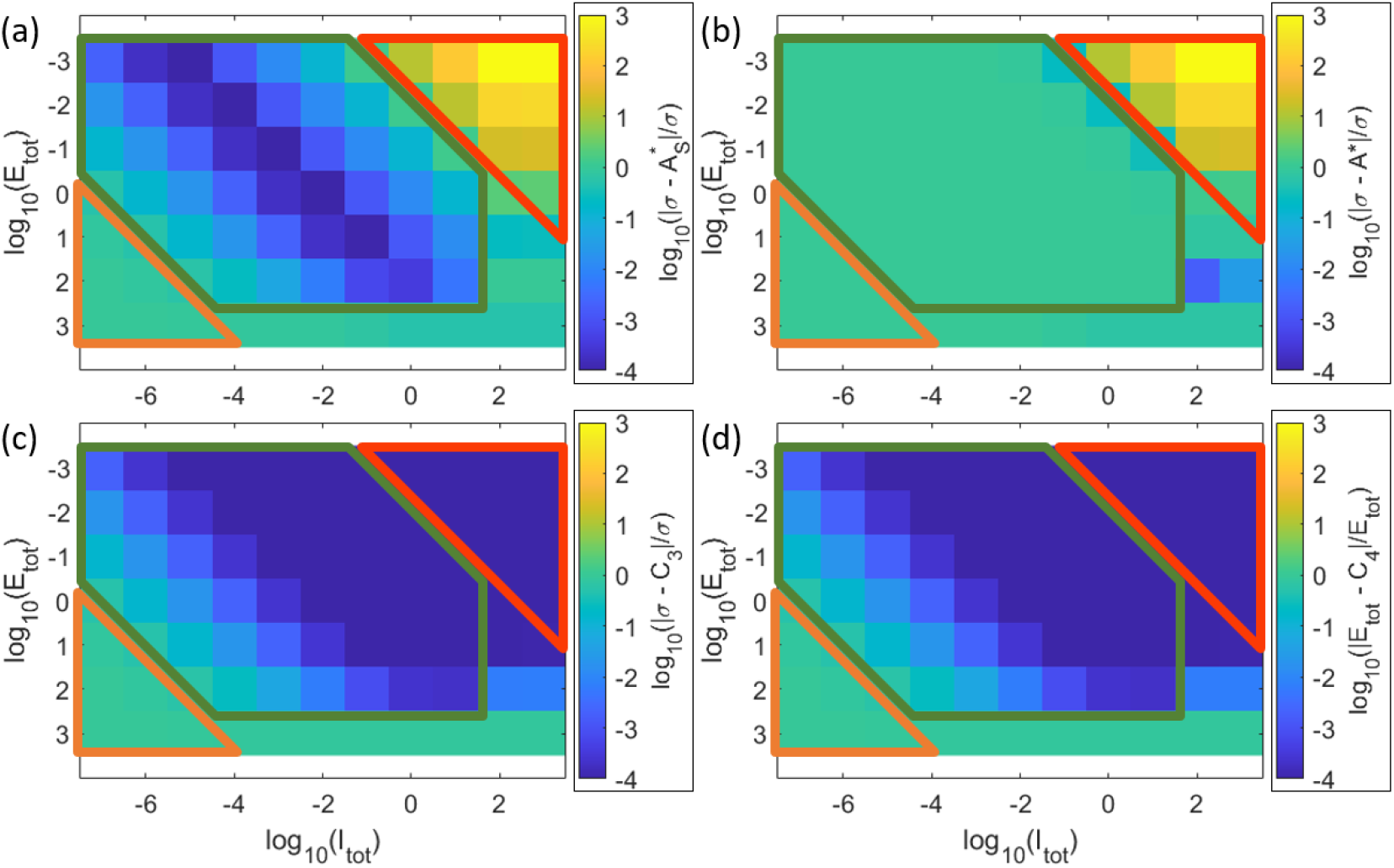
Comparison of protein concentrations for a wide range of input values (*I*_*tot*_) and total enzyme abundances (*E*_*tot*_), during three phases of the network response: prior to RPA (orange region), during RPA (green region), and after RPA has been lost (red region). We demonstrate the abundances (relative to *σ*) of: (a) the total input to the opposer cycle, 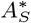; (b) the free, active output protein, *A*^*∗*^; (c) the complex *C*_3_; and (d) the complex *C*_4_ relative to the total enzyme abundance, *E*_*tot*_. Each heatmap represents the logged relative error of the proteins indicated in the label. As shown, RPA is associated with the conditions 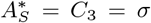, with *A*^*∗*^ = 0, and *C*_4_ = *E*_*tot*_. Parameters: *A*_*tot*_ = *B*_*tot*_ = 200, *k*_*A*1_ = *k*_*A*2_ = 200, *k*_*B*1_ = 10, *k*_*B*2_ = 4, *K*_*A*1_ = *K*_*A*2_ = 1, *K*_*B*1_ = *K*_*B*2_ = 0.01, *d*_*A*1_ = *d*_*A*2_ = *d*_*B*1_ = *d*_*B*2_ = 1, and 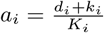.

In Fig 5, we provide a representative comparison between the dose-response profile of an RPA-exhibiting Michaelian model (as used by Ma et al. [8]) and that of the corresponding complex-complete model. As shown, the Michaelian model bears all the hallmarks of RPA: one variable (*A*^*∗*^) which maintains a fixed value (*σ*) over a range of system inputs, while another variable in the network (*B*^*∗*^) varies with the input. For the corresponding complex-complete model, on the other hand, *A*^*∗*^ increases monotonically with the system input, for all inputs; by contrast, *B*^*∗*^ initially increases steeply with the input, then gradually decreases, thereby exhibiting a prozone effect. (We refer the interested reader to [45] for other examples of covalent-modification cycles that exhibit a prozone effect of this type).

**Figure 5:**
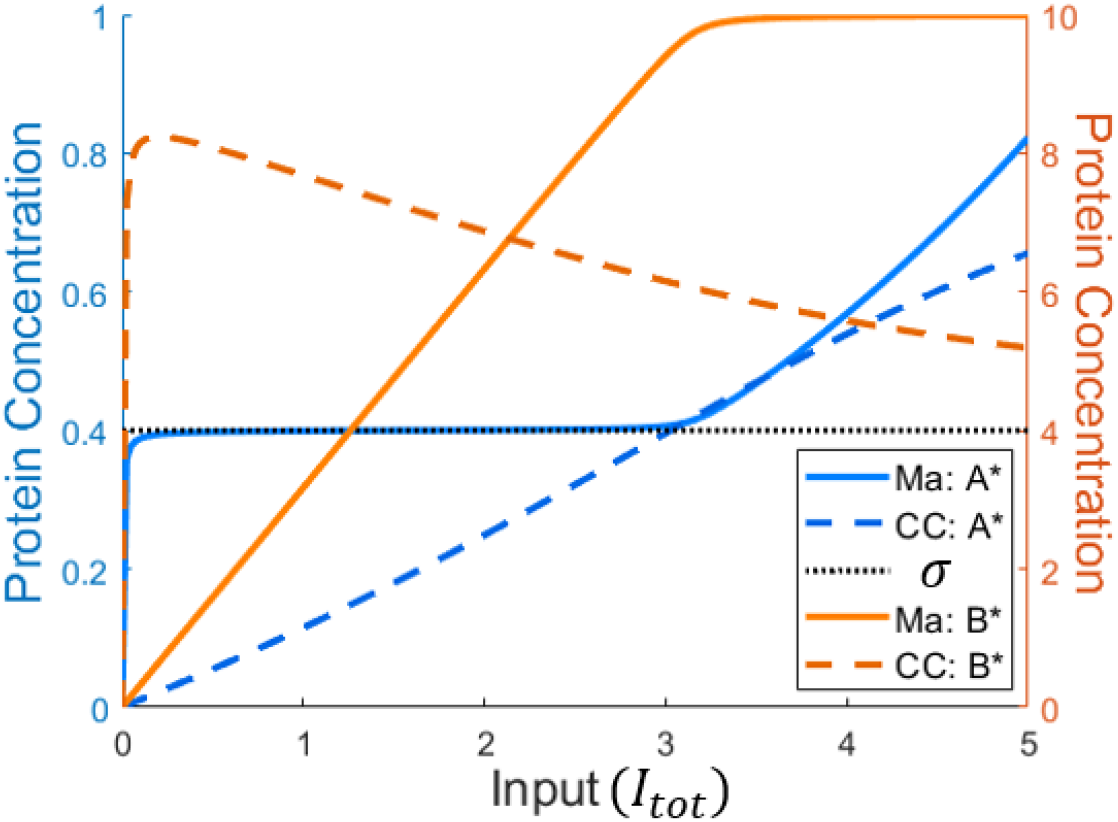
Free active protein forms in the Ma et al. (solid lines) [8] and complex-complete (CC) (dashed lines) models under the same parameter regime. Parameters: *A*_*tot*_ = *B*_*tot*_ = 10, *E*_*tot*_ = 1, *k*_*A*1_ = *k*_*A*2_ = 200, *k*_*B*1_ = 10, *k*_*B*2_ = 4, *K*_*A*1_ = *K*_*A*2_ = 1, *K*_*B*1_ = *K*_*B*2_ = 0.01, *d*_*A*1_ = *d*_*A*2_ = *d*_*B*1_ = *d*_*B*2_ = 1, and 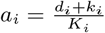.

When 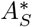 is considered, on the other hand, as depicted in Fig 6, we find that RPA is achieved, with 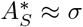 (and *A*^*∗*^ = 0), albeit for a very small range of inputs (Fig 6(a)). As the input to the system is increased by several orders of magnitude, however, 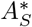 readily exceeds the specified tolerance around the estimated setpoint and increases monotonically with the system input. Strikingly, we see that RPA is lost precisely when *A*^*∗*^, the putative RPA variable in the Michaelian approximation of the system’s rate equations, acquires a non-zero value: the concentration of the complex *C*_3_ continues to track a value of *σ*, (Fig 6(b)) - an instance of ‘trivial’ RPA, since the output to the opposer cycle, 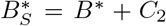 has reached its maximum value and no longer varies with the system input (Fig 6(b)).

**Figure 6:**
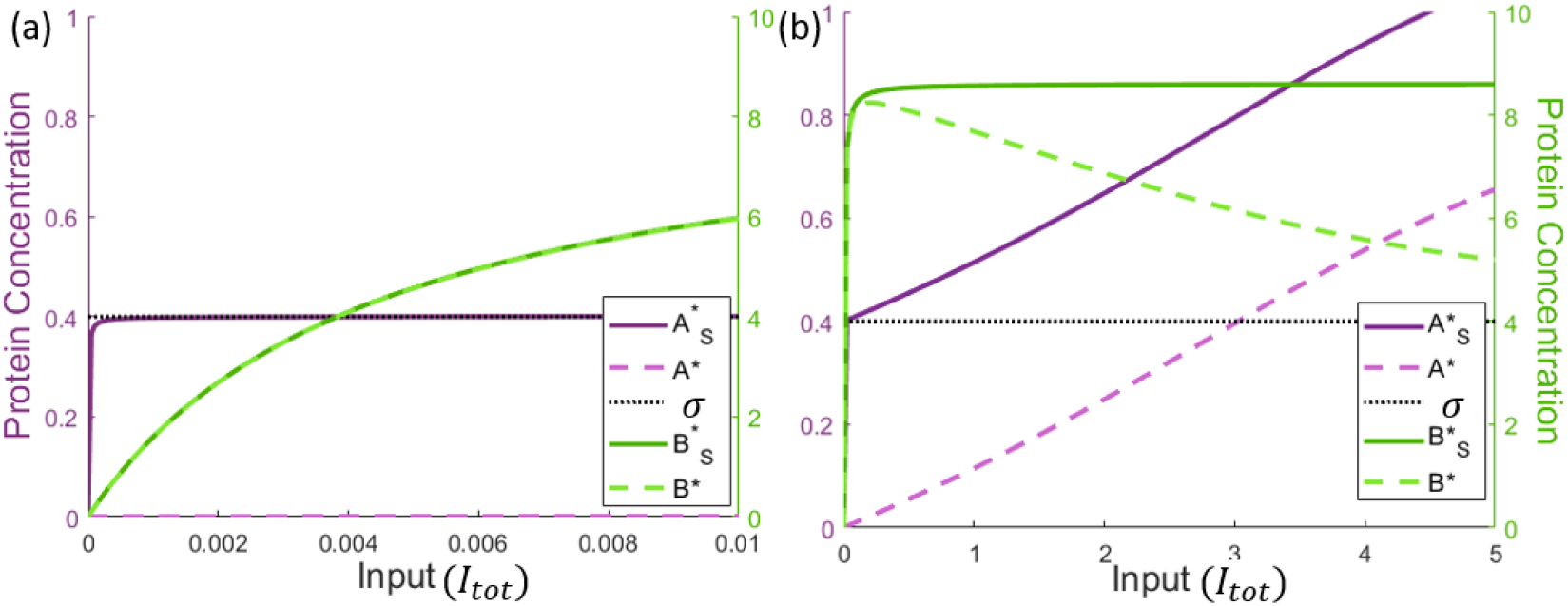
Comparison of the free active proteins (dashed lines) and the corresponding total active protein pools (comprising free protein plus complex with downstream substrate, solid lines) in the complex-complete model, for two input ranges: (a) *I*_*tot*_ ∈ [0, 10^−2^] and (b) *I*_*tot*_ ∈ [0, 5] (as seen in Fig 5). The total active protein pools are given by: 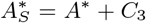 and 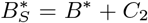. In (a) we restrict the input domain in order to demonstrate how the complex-complete network is capable of achieving RPA, whilst (b) demonstrates the deviation of the output from the estimated setpoint. Parameters: *A*_*tot*_ = *B*_*tot*_ = 10, *E*_*tot*_ = 1, *k*_*A*1_ = *k*_*A*2_ = 200, *k*_*B*1_ = 10, *k*_*B*2_ = 4, *K*_*A*1_ = *K*_*A*2_ = 1, *K*_*B*1_ = *K*_*B*2_ = 0.01, *d*_*A*1_ = *d*_*A*2_ = *d*_*B*1_ = *d*_*B*2_ = 1, and 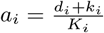.

### 3.2 Previously unrecognised constraints on protein abundances to support RPA

In our extensive simulations of the two versions of the two-protein pathway, both Michaelian and complex-complete, we found that the capacity for RPA is limited by the total abundances of the interacting proteins. Strikingly, the constraints on protein abundances required to support RPA are much more severe for the complex-complete model, than is predicted by the Michaelian simplification.

In Fig 7, we show that for the Michaelian simplification, RPA can only obtain (in the variable *A*^*∗*^) if 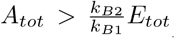 (Fig 7(a,c)), while no such constraint exists for the total abundance of the opposer protein (*B*_*tot*_) (Fig 7(b,d)). Nevertheless, we discovered that *B*_*tot*_ exerts a strong influence on the RPA range, independently of the value of *E*_*tot*_, once *E*_*tot*_ exceeds a threshold value determined by the sensitivity of the input/output cycle (governed by the values of *K*_*A*1_ and *K*_*A*2_), see Fig 7(b,d).

**Figure 7:**
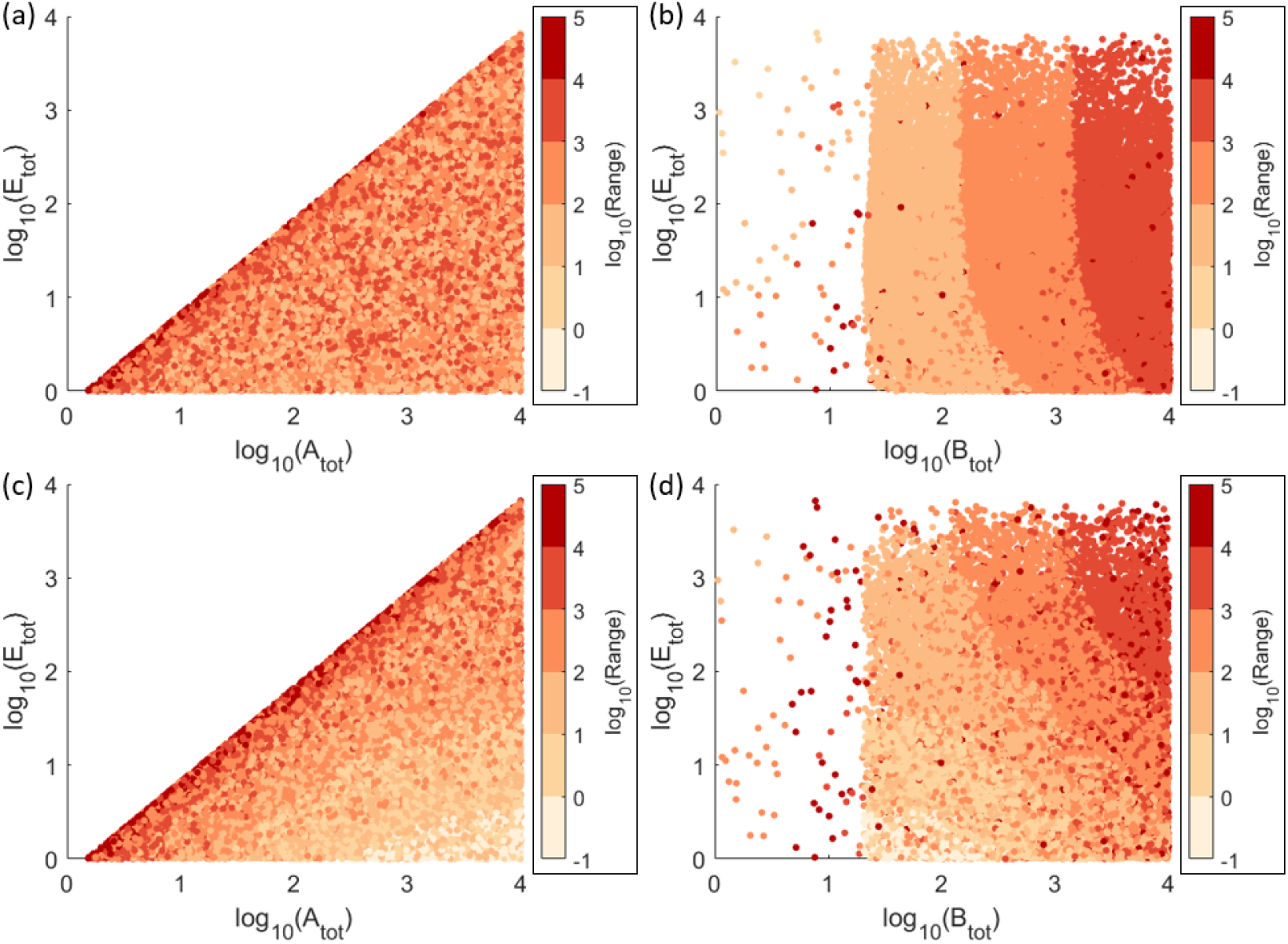
The influence of total substrate and enzyme abundances on RPA-capacity in the Michaelian (Ma et al. [8]) model for two sensitivity levels in the input/output cycle: (a) low sensitivity (*K*_*A*1_ = 700, *K*_*A*2_ = 500), (b) high sensitivity (*K*_*A*1_ = 7, *K*_*A*2_ = 5). When RPA obtains, the RPA Range is indicated by colour. As shown, the Michaelian model of RPA imposes strict constraints on *A*_*tot*_ relative to *E*_*tot*_ (a,c), with no constraints on *B*_*tot*_ (b,d). However *B*_*tot*_ exerts a significant influence on the RPA range. Parameters: *k*_*A*1_ = 7, *k*_*A*2_ = 5, *k*_*B*1_ = 2, *k*_*B*2_ = 3, *K*_*B*1_ = 2 ∗ 10^−2^, and *K*_*B*2_ = 3 ∗ 10^−2^.

For the complex-complete model, on the other hand, we find that while the constraint 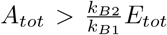 still holds (Fig 8(a,c)), the capacity for RPA also strictly requires 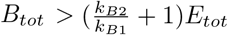 (Fig 8(b,d)). Moreover, both *B*_*tot*_ and *E*_*tot*_ exert a strong influence on RPA range (Fig 8(b,d)).

**Figure 8:**
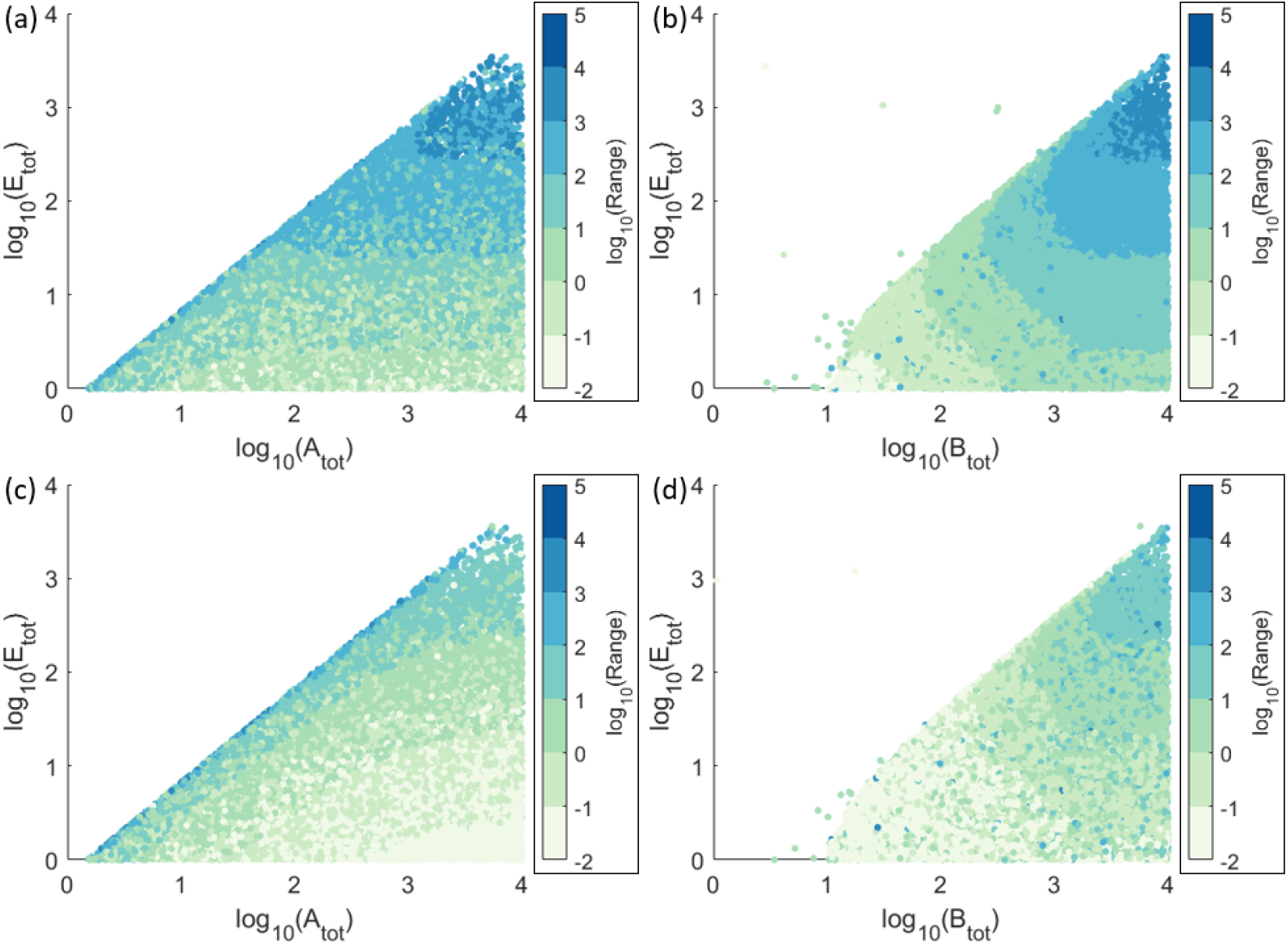
The influence of total substrate and enzyme abundances on RPA-capacity in the complex-complete model for two sensitivity in the input/output cycle: (a) low sensitivity (*K*_*A*1_ = 700, *K*_*A*2_ = 500), (b) high sensitivity (*K*_*A*1_ = 7, *K*_*A*2_ = 5). When RPA obtains, the RPA range is indicated by colour. As shown, the complex-complete model of RPA imposes strict constraints on all total protein abundances, all of which can also affect the RPA Range. Parameters: *k*_*A*1_ = 7, *k*_*A*2_ = 5, *k*_*B*1_ = 2, *k*_*B*2_ = 3, *K*_*A*1_ = 7, *K*_*A*2_ = 5, and *K*_*B*1_ = 2 10^−2^, *K*_*B*2_ = 3 10^−2^, *d*_*A*1_ = *d*_*A*2_ = *d*_*B*1_ = *d*_*B*2_ = 1, and 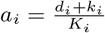.

That the condition 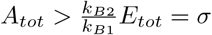 must be satisfied for RPA to obtain is self-evident, since the total abundance of the protein *A* must be at least as great as the setpoint. This condition must therefore hold for *any* model of RPA, however approximate or simplified. But when the presence of protein-protein complexes are taken into account, the protein abundance requirements of the network are more subtle. Indeed, an analysis of the system’s steady states, taking into consideration its mass conservation relations, makes clear that the presence of enzyme-substrate complexes imposes a number of additional constraints on the total protein abundances required for RPA.

Recall that *B*_*tot*_ = *B* + *B*^*∗*^ + *C*_2_ + *C*_3_ + *C*_4_ (see system equations given in Methods). We showed in Section 3.1 that when RPA obtains, 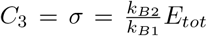 and *A*^*∗*^ = 0, with *C*_4_ = *E*_*tot*_ and *E*_1_ = 0. Under these conditions, the maximum value of 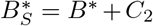 occurs when *B* = 0, giving

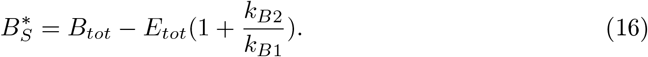

Now, considering the input/output cycle in isolation, detached from the opposer cycle (Fig 9(a)), the output for that cycle, 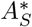, approaches its half-maximal value as 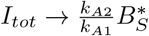(see [45, 62] for further details on this point). Therefore, if the maximum possible 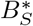 for the system is increased, a greater range of values for *I*_*tot*_ are compatible with achieving the RPA setpoint 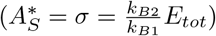 (see Fig 9(b)). Thus, for a given value of *E*_*tot*_, increasing *B*_*tot*_ increases the RPA range. Moreover, Equation 16 makes clear that the capacity for RPA strictly requires 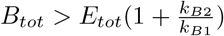.

In addition, since *E*_*tot*_ determines the setpoint, which is tracked by the variable 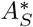 under RPA-permissive conditions, it is clear from the structure of the input/output cycle (see Fig 9(a)) that provided the minimum requirements for *A*_*tot*_ and *B*_*tot*_ are met, a greater range of *I*_*tot*_ values are compatible with RPA for a larger value of *E*_*tot*_ (with a proportionally larger value of 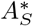) (see Fig 9(b)), thereby allowing a larger RPA range.

**Figure 9:**
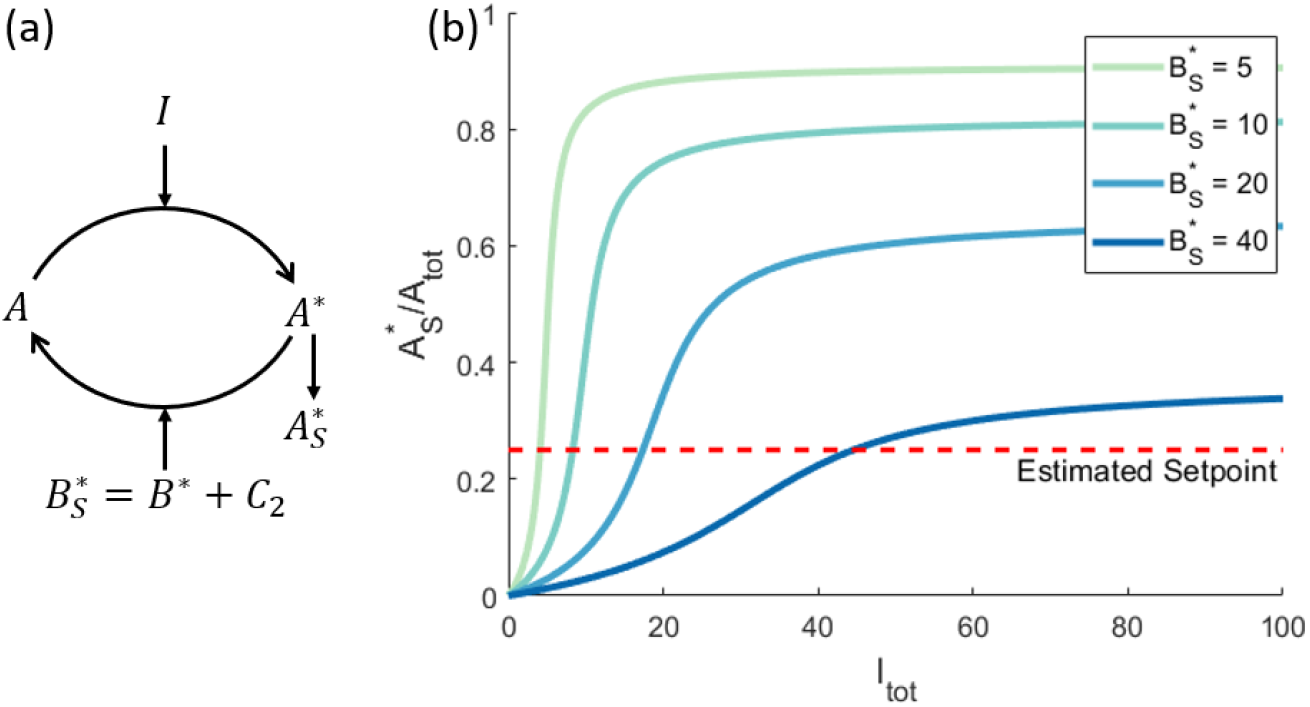
(a) The input/output cycle, considered as a single reversible covalent-modification cycle. (b) The dose response profile for this cycle, for varying enzyme abundances, 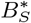. As shown, increasing 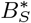 increases the amount of input (*I*_*tot*_) required to generate a particular output value for 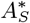. When considering the maximum possible value that 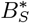 can take (as determined by the opposer cycle in the full system), the intersection of a curve with the estimated setpoint line indicates the maximum value of *I*_*tot*_ for which RPA obtains. If the maximum possible value of 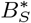 is reduced (lighter curves), the maximum possible value of *I*_*tot*_ for which RPA obtains is reduced commensurately, thereby reducing the RPA range. Parameters: *A*_*tot*_ = 100, *k*_*A*1_ = *k*_*A*2_ = *k*_*B*1_ = *k*_*B*2_ = 1, *K*_*A*1_ = *K*_*A*2_ = 0.01, *d*_*A*1_ = *d*_*A*2_ = 1, and 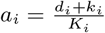.

Most significantly, our extensive computational searches, taken together, highlight the fact that the Michaelian model of ultrasensitivity-driven RPA characteristically overestimates the RPA range, in comparison with the ‘true’ RPA range calculated by the complex-complete framework. In fact, although the assumption of negligible enzyme-substrate complexes (central to the Michaelian simplification) requires *A*_*tot*_, *B*_*tot*_ ≫ *E*_*tot*_, i.e. substrate concentrations exist in vast excess over enzyme concentrations, we found that the greatest agreement between the two modelling frameworks occurred when *A*_*tot*_ ≈ *B*_*tot*_ ≈ *E*_*tot*_. Furthermore, the more the total protein concentrations *A*_*tot*_, *B*_*tot*_ exceed the enzyme concentration *E*_*tot*_, the more the Michaelian model overestimates the RPA range (Fig 10). Paradoxically, then, under the very conditions that might seem to justify the use of the Michaelis-Menten equation, this simplified model is most misleading in its predictions of the system’s RPA capacity.

**Figure 10:**
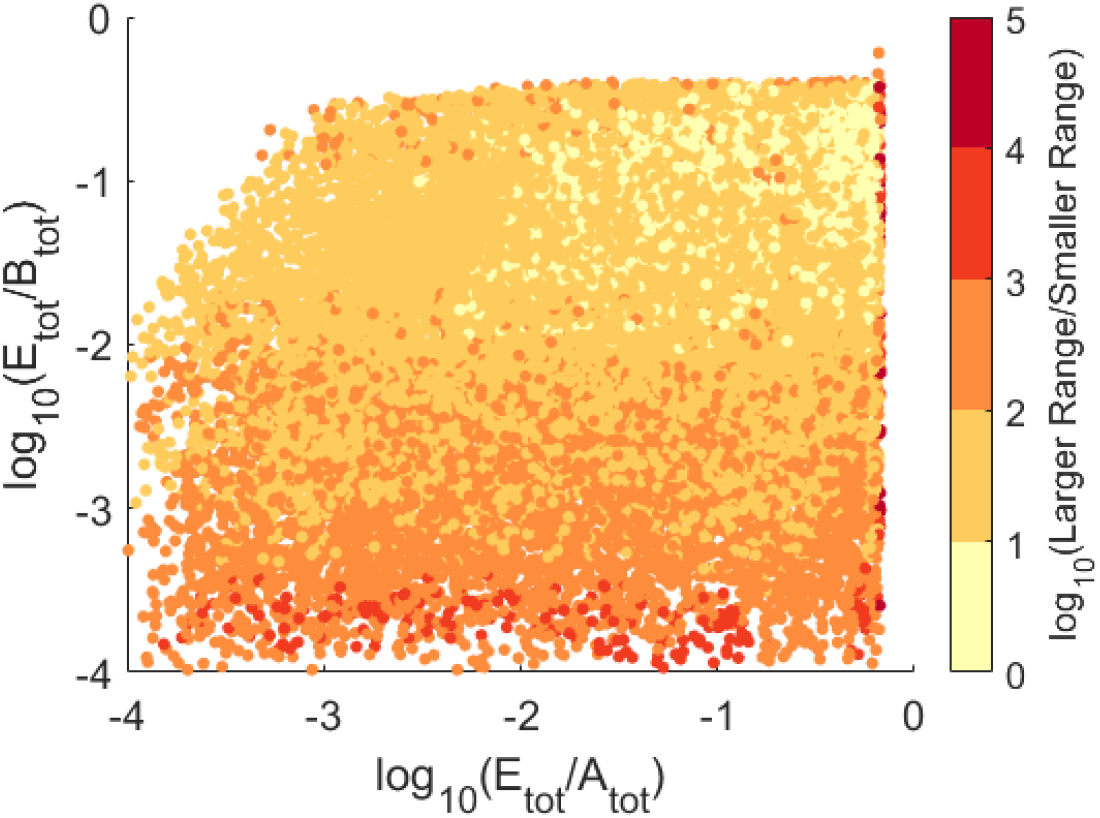
Increasing *E*_*tot*_ relative to *A*_*tot*_ and *B*_*tot*_ in both the Ma et al. [8] and complex-complete models results in the convergence of the RPA Ranges across the two model types. For each parameter choice, we compare the RPA Ranges predicted by the two different models, in each case dividing the larger range by the smaller range. Note that the RPA range for the Michaelian model is always the larger of the two. The log of this result is thus greater than zero, and the closer to zero (light yellow) the result becomes, the more similar are the RPA ranges for the two models. Parameters shown here: *k*_*A*1_ = 7, *k*_*A*2_ = 5, *k*_*B*1_ = 2, *k*_*B*2_ = 3, *K*_*A*1_ = 7, *K*_*A*2_ = 5, *K*_*B*1_ = 2 ∗ 10^−2^, *K*_*B*2_ = 3 ∗ 10^−2^, *E*_*tot*_, *A*_*tot*_, *B*_*tot*_ ∈ [1, 10^4^], *d*_*A*1_ = *d*_*A*2_ = *d*_*B*1_ = *d*_*B*2_ = 1, and 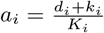.

### 3.3 RPA imposes parametric constraints on both Opposer and Input/Output cycles

In their simplified (Michaelian) model of RPA, neglecting the presence of protein-protein complexes, Ma et al. [8] identified a very specific parameter requirement for RPA - namely, very small Michaelis constants in the opposer cycle relative to the total abundance of the opposer protein. As noted earlier, this specific parametric condition engenders zero-order ultrasensitivity in the opposer cycle. Moreover, Ma et al. [8] showed analytically that, in the Michaelian framework, the presence of RPA was unaffected by any other parameters in the network.

By contrast, once all protein-protein interactions and intermediate complexes are included, we find that RPA imposes tight parameter constraints on *both* the opposer cycle *and* the input/output cycle. We confirm that RPA requires small Michaelis constants in the opposer cycle, *K*_*B*1_ and *K*_*B*2_ (Fig 11(a)), and that as these Michaelis constants increase (Fig 11(b)), the RPA property is lost. However, we also identified a significant region of our parameter space with very small values of *K*_*B*1_ and *K*_*B*2_, for which RPA did not obtain. On closer examination of parameter sets associated with RPA, we found that the Michaelis constants, *K*_*A*1_ and *K*_*A*2_, as well as the catalytic constants, *k*_*A*1_ and *k*_*A*2_, for the input/output cycle, had a significant impact on the ability of the network to exhibit RPA and on the RPA range.

**Figure 11:**
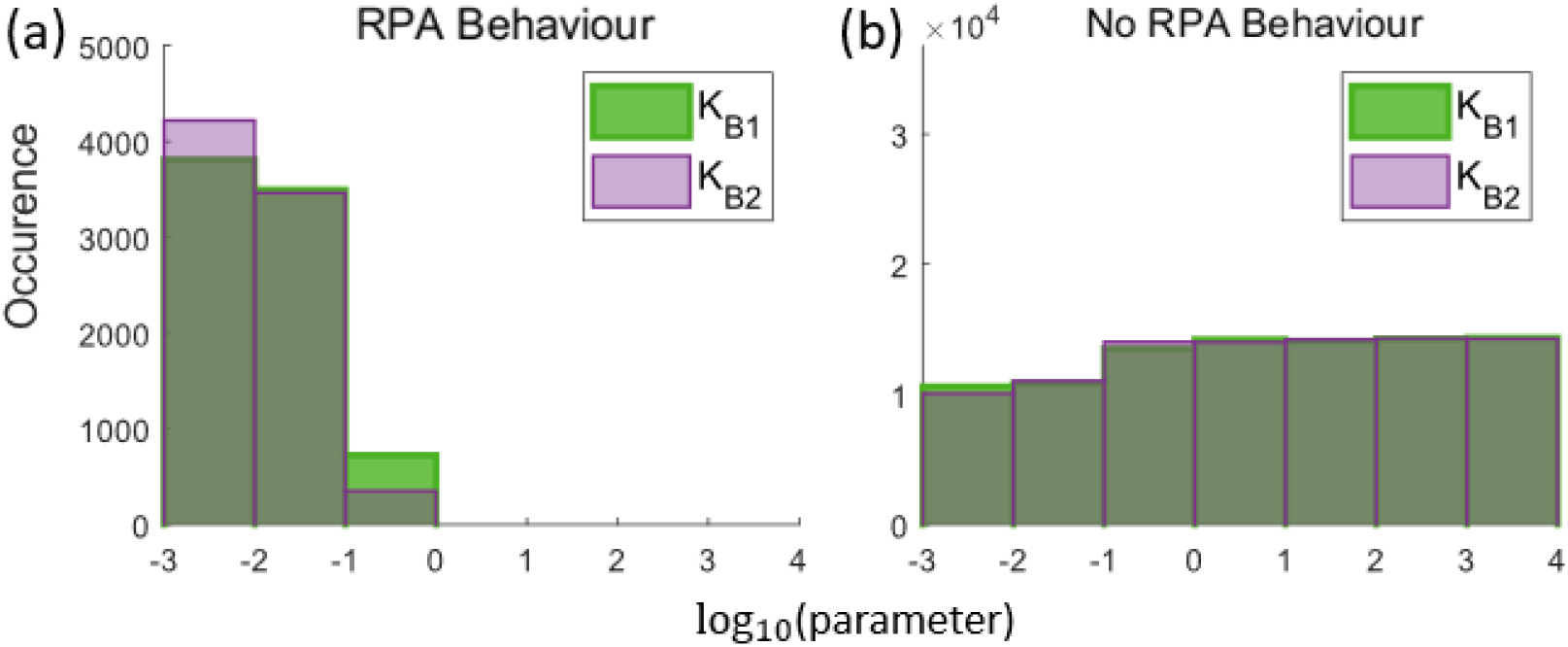
Histograms of (a) RPA vs (b) non-RPA responses for varying Michaelis constants in the opposer cycle. Small Michaelis constants are a necessary, but insufficient, condition for generating the RPA response. 10^5^ simulations were run with Michaelis constants being randomly selected from 10^−3^ to 10^4^. Parameters: *A*_*tot*_ = *B*_*tot*_ = 10, *E*_*tot*_ = 1, *k*_*A*1_ = *k*_*A*2_ = *k*_*B*1_ = *k*_*B*2_ = 1, *d*_*A*1_ = *d*_*A*2_ = *d*_*B*1_ = *d*_*B*2_ = 1, and 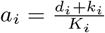.

Indeed, we found that RPA in the complex-complete framework requires large values of *K*_*A*1_, and small values of *K*_*A*2_. As we show in Fig 12a-c, decreasing the value of *K*_*A*1_ and increasing the value of *K*_*A*2_ reduces the RPA range. Likewise, RPA is promoted by small values of *k*_*A*1_ and large values of *k*_*A*2_, with a marked reduction in RPA range resulting from increasing values of *k*_*A*1_ and reducing values of *k*_*A*2_ (Fig 12d-f).

**Figure 12:**
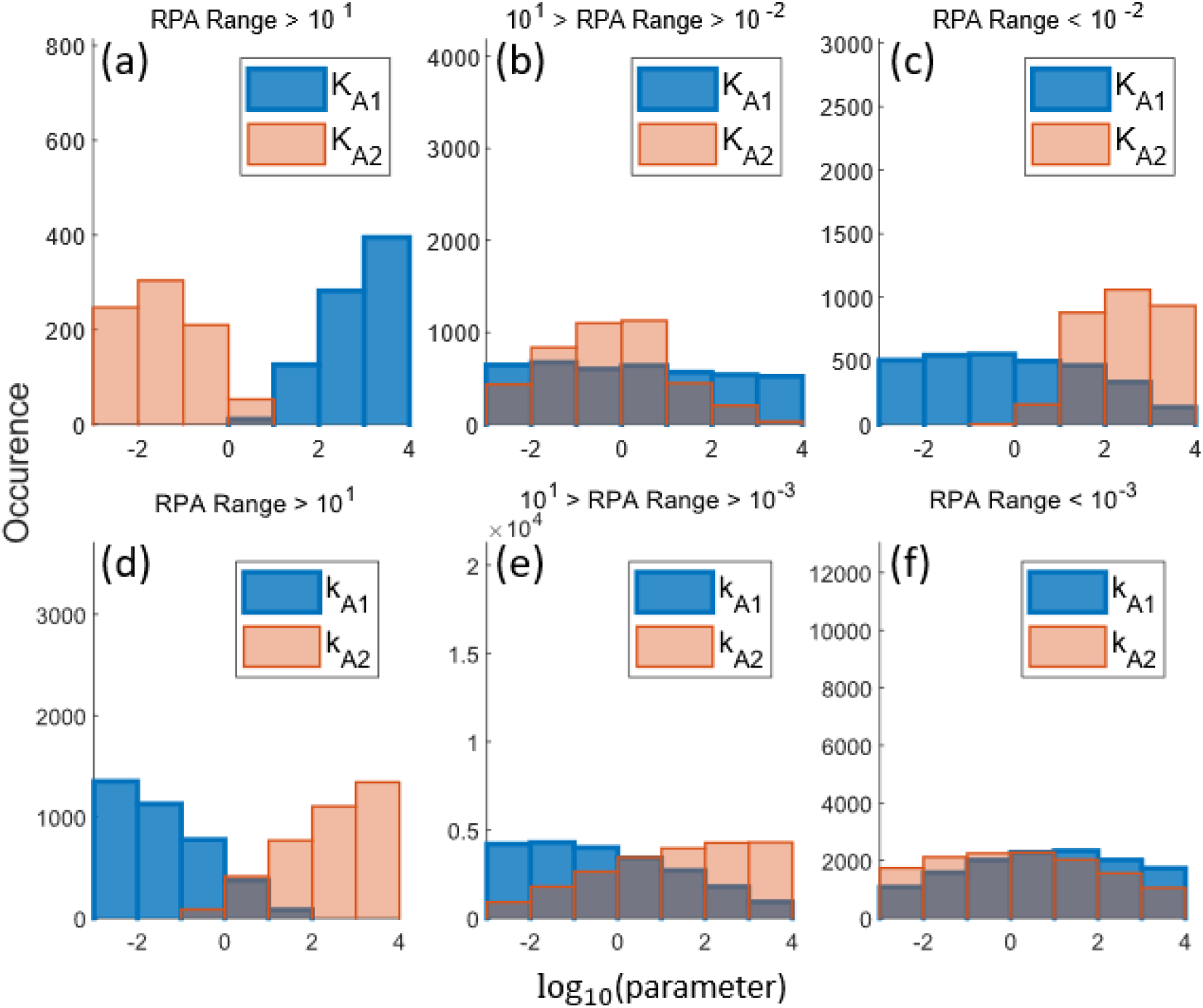
Histograms of the input/output cycle Michaelis constants (a-c), and catalytic constants (d-f), which were associated with RPA, grouped by the magnitude of the RPA Range. As shown, smaller values of *K*_*A*2_ and *k*_*A*1_, and larger values of *K*_*A*1_ and *k*_*A*2_ are associated with larger RPA Ranges. 10^5^ simulations were run with Michaelis constants or catalytic constants being randomly selected from 10^−3^ to 10^4^. Parameters: *A*_*tot*_ = *B*_*tot*_ = 10, *E*_*tot*_ = 1, *K*_*A*1_ = *K*_*A*2_ = 1 (except where noted otherwise), *K*_*B*1_ = *K*_*B*2_ = 0.01, *k*_*A*1_ = *k*_*A*2_ = *k*_*B*1_ = *k*_*B*2_ = 1 (except where noted otherwise), *d*_*A*1_ = *d*_*A*2_ = *d*_*B*1_ = *d*_*B*2_ = 1, and 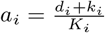.

The role and significance of these parameters may be elucidated by examining the input/output cycle as a single reversible covalent-modification cycle, as previously implemented in Fig 9. Applying either of the parameter conditions which increase the RPA range, for the Michaelis (Fig 13(a)) or the catalytic constants (Fig 13(b)), results in dose-response profiles which require more input to achieve equivalent output concentrations. Importantly, the dose-responses in Fig 13 are underpinned by increased protein-protein complex concentrations, and cannot be obtained by a simplified Michaelian framework, as used by Ma et al. [8].

**Figure 13:**
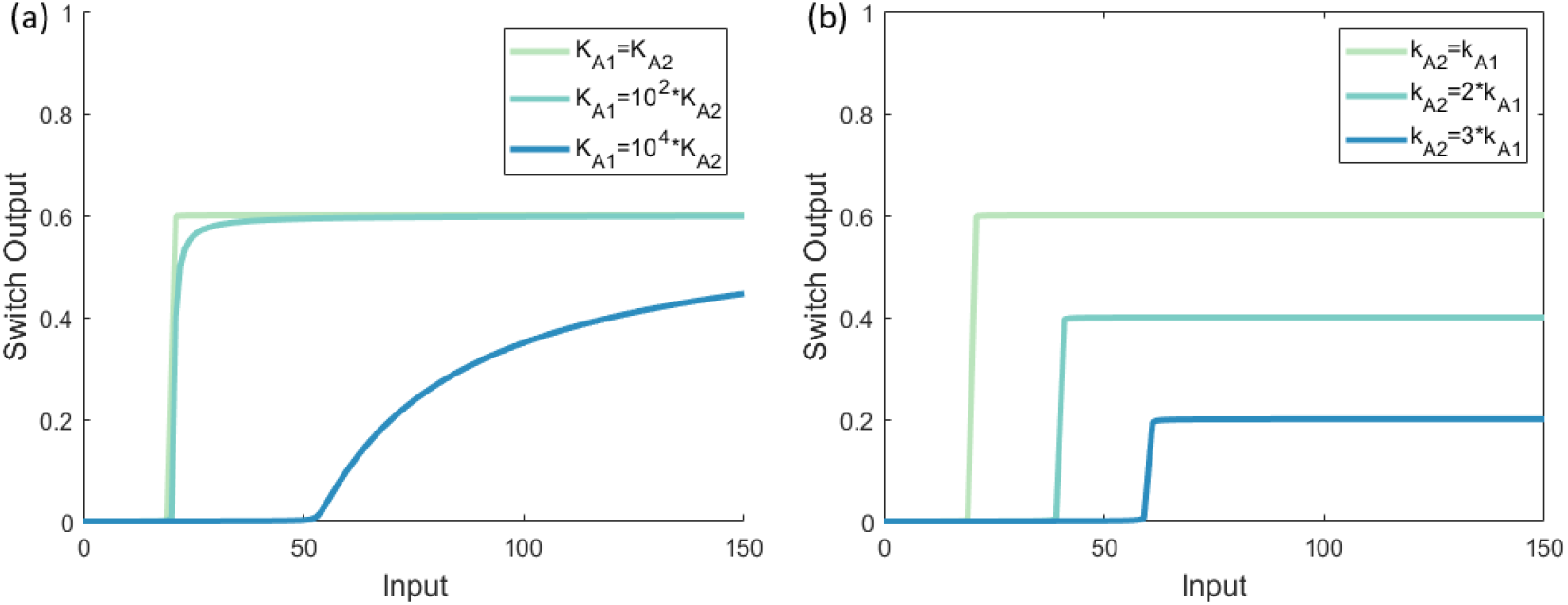
The role of (a) Michaelis constants and (b) catalytic constants in a single reversible covalent-modification cycle. Parameters: *A*_*tot*_ = 100, *E*_*tot*_ = 20, *k*_*A*1_ = *k*_*A*2_ = 1, *K*_*A*1_ = *K*_*A*2_ = 0.01, *d*_*A*1_ = *d*_*A*2_ = 1, and 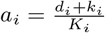.

## 4 Discussion and Concluding Remarks

Ma et al. [8] identified the earliest known instances of what are now known as Opposer Modules and Balancer Modules, generalisations of which were later shown definitively to be topological basis modules for all possible RPA-capable networks of any size and complexity [4]. Both classes of RPA basis module exhibit strict structural requirements (involving an overarching feedback structure in the case of Opposer modules, and a feedforward structure in the case of Balancer modules) and, crucially, execute their RPA-conferring computational functions through the embedding of certain types of computational ‘nodes’ into these well-defined structures – opposer nodes, embedded into the feedback component(s) of Opposer Modules; and balancer nodes, embedded into the feedforward component(s) of Balancer Modules.

Unlike Balancer nodes, which are relatively easy to construct [40], opposer nodes are now acknowledged to be notoriously difficult to create at the level of intermolecular interactions, in the form of chemical reaction networks (CRNs) [39]. The Ma et al. study [8] used the commonly-employed simplification of Michaelis-Menten kinetics to model three-node networks of interlinked covalent-modification cycles, and used this modelling approximation to suggest that an ultrasensitivity-generating mechanism suffices as an opposer node (or a ‘buffer node’ in the language of that early study). As a consequence, it has been taken for granted by the life-sciences community that RPA can readily be orchestrated in signalling networks constructed from collections of enzyme-mediated post-translational modifications.

Here we show that, when the detailed intermolecular interactions required for enzyme-mediated reactions are fully accounted for, RPA is much more difficult to achieve via an ultrasensitivity-generating mechanism than previously thought. In fact, RPA cannot be achieved, under any parametric conditions, in the ‘free’ activated form of any protein – a remarkable and functionally crucial result that has been overlooked in all previous studies of RPA. Strikingly, the *only* species that can possibly achieve RPA for a non-negligible range of disturbances to the network is the ‘total’ active pool of each protein that resides, topologically speaking, between the opposer node (which exhibits ultrasensitivity) and the ‘connector node’ of the Opposer module (see [4, 39] for a comprehensive overview of Opposer module topologies). This total active protein pool comprises both the free form of the activated protein, as well as a complexed form with its downstream protein substrate. In the case of the protein which directly regulates the opposer node (the latter often being referred to in the language of control theory as the *sensor* node), this total pool of active protein comprises *only* the enzyme-substrate complex, with zero concentration in the free form. This particular protein thereby exhibits RPA in the concentration of a protein-protein complex that is neglected entirely in the Michaelian framework.

For our model, being a minimal Opposer module, the sensor protein (A) and the opposer protein (B) are the only two proteins considered, with the sensor protein also playing the additional role of input/output node for the module. It is clear from the definitive topological properties of Opposer modules that any additional proteins between the sensor and the connector nodes (see [4]) will also exhibit RPA; from our complex-complete model reported here, it follows that each of these proteins will exhibit RPA in the total pool of the activated form, but will generally comprise a non-zero concentration of the free form since these proteins regulate a *non-ultrasensitive* downstream protein ([45]). Indeed, the partition of the RPA-exhibiting total pool into free and complexed forms depends on the sensitivity of the protein immediately downstream, and will therefore be highly variable ([45]).

In any case, since RPA can only obtain in a protein pool that comprises both a free and a complexed form of a protein, the biological usefulness of this form of RPA is dependent upon the ability of such a protein to orchestrate its downstream extramodular functions equally in either form. In other words, the conformation of an RPA-exhibiting protein must allow it to bind its downstream target(s) with equal affinity in either its free or complexed (with its intramodular substrate) form. Should ultrasensitivity-dependent RPA actually be implemented by complex cellular networks, this necessary structural property of RPA-capable proteins may provide valuable clues as to the portion of the proteome that could participate in such RPA-conferring networks.

Beyond the identity of the RPA-capable network variable, our complex-complete modelling framework also highlights the fact that RPA places much greater constraints on the feasible parameter space than suggested by the Michaelian framework [8]. In addition, our work also considers the important concept of ‘RPA range’ – that is, the range of network disturbances or stimuli for which RPA obtains – which has largely been overlooked in previous studies, which have focussed purely on the presence or absence of RPA. This is a crucially important property, since networks with only a vanishingly small RPA range may provide no functional benefits, in practice, in comparison with networks that cannot exhibit RPA at all. In this connection, it is intriguing to observe that parameter alterations that increase RPA range could ultimately jeopardize the very presence of RPA. For instance, we have shown that decreasing the abundances of protein substrates (e.g. *A*_*tot*_) tends to increase the RPA range; but we also show that RPA is only possible for protein abundances above a certain threshold 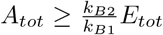. In a population of cells, where protein abundances are inevitably highly variable, it may thus be advantageous to the robustness of signalling functions (e.g. via RPA capacity) across the entire population to limit the total expression levels of proteins to maintain enzymes and substrates at comparable concentrations; on the other hand, this strategy risks the loss of robustness altogether in a subset of the population should substrate abundances in those cells fall below the threshold level.

Importantly, our study highlights the fact that Michaelian models characteristically over-estimate the RPA range. In fact, the very conditions which are generally thought to justify the use of the Michaelian simplification (protein substrate concentrations, e.g. *A*_*tot*_, *B*_*tot*_, in vast excess over enzyme concentrations, e.g. *E*_*tot*_), result in the largest discrepancy in RPA range in comparison with the complex-complete framework.

The shortcomings of the Michaelis-Menten equation to accurately capture the processing of biochemical signals via enzyme-mediated signalling events are now widely acknowledged, particularly in contexts involving highly intricate intermolecular interactions, and this study adds to the growing body of literature that cautions against its indiscriminate use [45, 46, 49, 57, 58, 50, 51, 52, 53, 54]. Michaelian models of RPA such as those developed by Ma et al. [8] assume *a priori* that protein-protein complexes exist at sufficiently small concentrations that they can be neglected. By contrast, the present study makes clear that the existence of protein-protein complexes is key to the very existence of RPA via ultrasensitivity-generating mechanisms, and *cannot* be neglected. The centrally important role of protein-protein complexes (e.g. in the form of sigma/anti-sigma factor complexes) in antithetic-integral control – the other type of opposer mechanism, distinct from the ultrasensitivity-dependent mechanism considered here – is already well-established [2]. Since RPA is a ubiquitously-observed and functionally crucial network response for biologic systems across all domains of life, the problem of constructing biologically-useful opposer nodes through intermolecular interactions, particularly in the context of vast and highly complex signal transduction networks where Opposer modules are considered most prevalent, is vitally important to the evolution of life.

## Data Accessibility

All code used in this study is available at: https://github.com/JeynesSmith/JeynesSmithPerfectAdaptation2022

## Author Contributions

CJS: conceptualisation, formal analysis, methodology, software, visualisation, writing - original draft preparation. RPA: conceptualisation, methodology, writing - review and editing, supervision.

## Acknowledgements

Computational resources and services used in this work were provided by the eResearch Office, Queensland University of Technology, Brisbane, Australia.

## References

[1] Ferrell Jr JE. 2016 Perfect and near-perfect adaptation in cell signaling. Cell systems 2, 62–67.

[2] Khammash MH. 2021 Perfect adaptation in biology. Cell Systems 12, 509–521.

[3] Araujo RP, Vittadello ST, Stumpf MP. 2021 Bayesian and Algebraic Strategies to Design in Synthetic Biology. Proceedings of the IEEE.

[4] Araujo RP, Liotta LA. 2018 The topological requirements for robust perfect adaptation in networks of any size. Nature communications 9, 1–12.

[5] Yi TM, Huang Y, Simon MI, Doyle J. 2000 Robust perfect adaptation in bacterial chemotaxis through integral feedback control. Proceedings of the National Academy of Sciences 97, 4649–4653.

[6] Araujo RP, Liotta LA, Petricoin EF. 2007 Proteins, drug targets and the mechanisms they control: the simple truth about complex networks. Nature reviews Drug discovery 6, 871–880.

[7] Araujo RP, Liotta LA. 2006 A control theoretic paradigm for cell signaling networks: a simple complexity for a sensitive robustness. Current opinion in chemical biology 10, 81–87.

[8] Ma W, Trusina A, El-Samad H, Lim WA, Tang C. 2009 Defining network topologies that can achieve biochemical adaptation. Cell 138, 760–773.

[9] Berg HC, Brown DA. 1972 Chemotaxis in Escherichia coli analysed by three-dimensional tracking. Nature 239, 500–504.

[10] Alon U, Surette MG, Barkai N, Leibler S. 1999 Robustness in bacterial chemotaxis. Nature 397, 168–171.

[11] Macnab RM, Koshland DE. 1972 The gradient-sensing mechanism in bacterial chemotaxis. Proceedings of the National Academy of Sciences 69, 2509–2512.

[12] Kirsch ML, Peters PD, Hanlon DW, Kirby JR, Ordal G. 1993 Chemotactic methylesterase promotes adaptation to high concentrations of attractant in Bacillus subtilis.. Journal of Biological Chemistry 268, 18610–18616.

[13] Mello BA, Tu Y. 2003 Quantitative modeling of sensitivity in bacterial chemotaxis: the role of coupling among different chemoreceptor species. Proceedings of the National Academy of Sciences 100, 8223–8228.

[14] Rao CV, Kirby JR, Arkin AP, Kirschner MW. 2004 Design and diversity in bacterial chemotaxis: a comparative study in Escherichia coli and Bacillus subtilis. PLoS biology 2, e49.

[15] Kollmann M, Løvdok L, Bartholomé K, Timmer J, Sourjik V. 2005 Design principles of a bacterial signalling network. Nature 438, 504–507.

[16] Endres RG, Wingreen NS. 2006 Precise adaptation in bacterial chemotaxis through “assistance neighborhoods”. Proceedings of the National Academy of Sciences 103, 13040–13044.

[17] Parent CA, Devreotes PN. 1999 A cell’s sense of direction. Science 284, 765–770.

[18] Yang L, Iglesias PA. 2006 Positive feedback may cause the biphasic response observed in the chemoattractant-induced response of Dictyostelium cells. Systems & control letters 55, 329–337.

[19] Levchenko A, Iglesias PA. 2002 Models of eukaryotic gradient sensing: application to chemotaxis of amoebae and neutrophils. Biophysical journal 82, 50–63.

[20] Barkai N, Leibler S. 1997 Robustness in simple biochemical networks. Nature 387, 913–917.

[21] Frei T, Chang CH, Filo M, Arampatzis A, Khammash M. 2022 A genetic mammalian proportional–integral feedback control circuit for robust and precise gene regulation. Proceedings of the National Academy of Sciences 119, e2122132119.

[22] Reisert J, Matthews HR. 2001 Response properties of isolated mouse olfactory receptor cells. The Journal of physiology 530, 113–122.

[23] Matthews HR, Reisert J. 2003 Calcium, the two-faced messenger of olfactory transduction and adaptation. Current opinion in neurobiology 13, 469–475.

[24] Hoeller O, Gong D, Weiner OD. 2014 How to understand and outwit adaptation. Developmental cell 28, 607–616.

[25] Kaupp UB. 2010 Olfactory signalling in vertebrates and insects: differences and commonalities. Nature Reviews Neuroscience 11, 188–200.

[26] Yau KW, Hardie RC. 2009 Phototransduction motifs and variations. Cell 139, 246–264.

[27] Hart Y, Madar D, Yuan J, Bren A, Mayo AE, Rabinowitz JD, Alon U. 2011 Robust control of nitrogen assimilation by a bifunctional enzyme in E. coli. Molecular cell 41, 117–127.

[28] Muzzey D, Gómez-Uribe CA, Mettetal JT, van Oudenaarden A. 2009 A systems-level analysis of perfect adaptation in yeast osmoregulation. Cell 138, 160–171.

[29] Shinar G, Feinberg M. 2010 Structural sources of robustness in biochemical reaction networks. Science 327, 1389–1391.

[30] El-Samad H, Goff J, Khammash M. 2002 Calcium homeostasis and parturient hypocalcemia: an integral feedback perspective. Journal of theoretical biology 214, 17–29.

[31] Ben-Zvi D, Barkai N. 2010 Scaling of morphogen gradients by an expansion-repression integral feedback control. Proceedings of the National Academy of Sciences 107, 6924–6929.

[32] Eldar A, Dorfman R, Weiss D, Ashe H, Shilo BZ, Barkai N. 2002 Robustness of the BMP morphogen gradient in Drosophila embryonic patterning. Nature 419, 304–308.

[33] Yadid G, Overstreet DH, Zangen A. 2001 Limbic dopaminergic adaptation to a stressful stimulus in a rat model of depression. Brain research 896, 43–47.

[34] Medina-Gomez G, Yetukuri L, Velagapudi V, Campbell M, Blount M, Jimenez-Linan M, Ros M, Orešič M, Vidal-Puig A. 2009 Adaptation and failure of pancreatic β cells in murine models with different degrees of metabolic syndrome. Disease models & mechanisms 2, 582–592.

[35] Sturgeon JA, Zautra AJ. 2010 Resilience: a new paradigm for adaptation to chronic pain. Current pain and headache reports 14, 105–112.

[36] Fodale V, Pierobon M, Liotta L, Petricoin E. 2011 Mechanism of cell adaptation: when and how do cancer cells develop chemoresistance?. Cancer journal (Sudbury, Mass.) 17, 89.

[37] Araujo R, Petricoin E, Liotta L. 2007 Mathematical modeling of the cancer cell’s control circuitry: paving the way to individualized therapeutic strategies. Current Signal Transduction Therapy 2, 145–155.

[38] Geho DH, Petricoin E, Liotta LA, Araujo R. 2005 Modeling of protein signaling networks in clinical proteomics. In Cold Spring Harbor symposia on quantitative biology vol. 70 pp. 517–524. Cold Spring Harbor Laboratory Press.

[39] Araujo RP, Liotta LA. 2020 Design Principles Underlying Robust Perfect Adaptation of Complex Biochemical Networks. bioRxiv.

[40] Araujo R, Liotta L. 2022 Universal structures for embedded integral control in biological adaptation. Research Square [https://doi.org/10.21203/rs.3.rs-1571178/v1].

[41] Goentoro L, Shoval O, Kirschner MW, Alon U. 2009 The incoherent feedforward loop can provide fold-change detection in gene regulation. Molecular cell 36, 894–899.

[42] Mangan S, Alon U. 2003 Structure and function of the feed-forward loop network motif. Proceedings of the National Academy of Sciences 100, 11980–11985.

[43] Briat C, Gupta A, Khammash M. 2016 Antithetic integral feedback ensures robust perfect adaptation in noisy biomolecular networks. Cell systems 2, 15–26.

[44] Aoki SK, Lillacci G, Gupta A, Baumschlager A, Schweingruber D, Khammash M. 2019 A universal biomolecular integral feedback controller for robust perfect adaptation. Nature 570, 533–537.

[45] Jeynes-Smith C, Araujo RP. 2021 Ultrasensitivity and bistability in covalent-modification cycles with positive autoregulation. Proceedings of the Royal Society A 477, e20210069.

[46] Kim JK, Tyson JJ. 2020 Misuse of the Michaelis–Menten rate law for protein interaction networks and its remedy. PLoS Computational Biology 16, e1008258.

[47] Segel LA, Slemrod M. 1989 The quasi-steady-state assumption: a case study in perturbation. SIAM review 31, 446–477.

[48] Goldbeter A, Koshland DE. 1981 An amplified sensitivity arising from covalent modification in biological systems. Proceedings of the National Academy of Sciences 78, 6840–6844.

[49] Gunawardena J. 2014 Time-scale separation–Michaelis and Menten’s old idea, still bearing fruit. The FEBS journal 281, 473–488.

[50] Ciliberto A, Capuani F, Tyson JJ. 2007 Modeling networks of coupled enzymatic reactions using the total quasi-steady state approximation. PLoS computational biology 3, e45.

[51] Fujioka A, Terai K, Itoh RE, Aoki K, Nakamura T, Kuroda S, Nishida E, Matsuda M. 2006 Dynamics of the Ras/ERK MAPK cascade as monitored by fluorescent probes. Journal of Biological Chemistry 281, 8917–8926.

[52] Blüthgen N, Bruggeman FJ, Legewie S, Herzel H, Westerhoff HV, Kholodenko BN. 2006 Effects of sequestration on signal transduction cascades. The FEBS journal 273, 895–906.

[53] Schnell S, Maini P. 2003 A century of enzyme kinetics: reliability of the KM and vmax estimates. Comments on theoretical biology 8, 169–187.

[54] Cha S. 1970 Kinetic behavior at high enzyme concentrations: magnitude of errors of Michaelis-Menten and other approximations. Journal of Biological Chemistry 245, 4814–4818.

[55] Tzafriri AR. 2003 Michaelis-Menten kinetics at high enzyme concentrations. Bulletin of mathematical biology 65, 1111–1129.

[56] Patsatzis DG, Goussis DA. 2019 A new Michaelis-Menten equation valid everywhere multi-scale dynamics prevails. Mathematical biosciences 315, 108220.

[57] Shinar G, Rabinowitz JD, Alon U. 2009 Robustness in glyoxylate bypass regulation. PLoS computational biology 5, e1000297.

[58] Xing J, Chen J. 2008 The Goldbeter-Koshland switch in the first-order region and its response to dynamic disorder. PloS one 3, e2140.

[59] Omale D, Ojih P, Ogwo M. 2014 Mathematical analysis of stiff and non-stiff initial value problems of ordinary differential equation using MATLAB. International journal of scientific & engineering research 5, 49–59.

[60] Aliyu B, Osheku C, Funmilayo A, Musa J. 2014 Identifying stiff ordinary differential equations and problem solving environments (PSEs). Journal of Scientific Research & Reports 3, 1430–1448.

[61] Ashino R, Nagase M, Vaillancourt R. 2000 Behind and beyond the MATLAB ODE suite. Computers & Mathematics with Applications 40, 491–512.

[62] Gomez-Uribe C, Verghese GC, Mirny LA. 2007 Operating regimes of signaling cycles: statics, dynamics, and noise filtering. PLoS computational biology 3, e246.

